# The immune response to a fungus in pancreatic cancer samples

**DOI:** 10.1101/2023.03.28.534606

**Authors:** KJ Brayer, JA Hanson, S Cingam, C Martinez, SA Ness, I Rabinowitz

## Abstract

Pancreatic ductal adenocarcinoma (PDAC) is a poor prognosis cancer with an .aggressive growth profile that is often diagnosed at late stage and that has few curative or therapeutic options. PDAC growth has been linked to alterations in the pancreas microbiome, which could include the presence of the fungus *Malassezia*. We used RNA-sequencing to compare 14 paired tumor and normal (tumor adjacent) pancreatic cancer samples and found *Malassezia* RNA in both the PDAC and normal tissues. Although the presence of *Malassezia* was not correlated with tumor growth, a set of immune- and inflammatory-related genes were up-regulated in the PDAC compared to the normal samples, suggesting that they are involved in tumor progression. Gene set enrichment analysis suggests that activation of the complement cascade pathway and inflammation could be involved in pro PDAC growth.

## Introduction

Pancreatic cancer (PDAC) is the 9^th^ most common cancer in the US but is the 4^th^ most common cause of cancer related death (∼54,000/year and ∼44,000/year respectively). The median 5 year survival for stage 4 disease is 9 % (1). The high death rate with respect to the prevalence rate is due to poor early detection and a lack of meaningful advancement in systemic therapeutics. The most common somatic mutations, Kirsten rat sarcoma viral oncogene (*KRAS*), tumor protein p53 (*TP53)*, cyclin dependent kinase inhibitor 2 A (*CDKN2A*), and SMAD family member 4 (*SMAD4*) have been identified by whole-exome and -genome sequencing of large PDAC cohorts and form the majority of unique mutations in patients with PDAC (2).

The tumor microenvironment (TME), which represents a complex ecosystem involving interactions between immune cells, cancer cells, stromal cells, and the extracellular matrix, can support tumor proliferation, survival, and metastasis and can be highly immunosuppressive (3,4,5).

The progression of PDAC growth requires micro environmental immune-suppressive inflammation in cooperation with oncogenic mutations. An association of the tumor bacterial microbiome in PDAC patients with short-term survival (STS) and long-term survival (LTS) was analyzed. An intra-tumoral microbiome signature (Pseudoxanthomonas, Streptomyces, Saccharopolyspora and Bacillus clausii) was highly predictive of LTS in both discovery and validation cohorts. They reproduced these results in a mouse model, by showing that the transfer of LTS and STS gut microbiomes altered the tumor microbiome and tumor growth in the mice.

(6) On the other hand, bacterial ablation in the pancreas was shown to alter the immune system, allowing the immune system response to decrease growth of the tumor by a reduction in myeloid-derived suppressor cells and an increase in M1 macrophage differentiation, promoting Th1 differentiation of CD4+ T cells and CD8+ T cell activation. Bacterial ablation also enabled efficacy for checkpoint-targeted immunotherapy by up-regulating PD-1 expression. (7) More recently the same group looked at the role of the fungal microbiome in PDAC. (8) In a mouse model they showed a significant migration of fungi from the lumen of the gut into the lumen of the pancreas in both mice and humans with PDAC compared to non-malignant controls. Both in mice and humans, the *Malassezia* fungus was highly enriched in the malignant pancreatic tumor. They demonstrated that the binding of mannose-binding lectin (MBL) to glycans of the fungal wall, and lectin pathway activation was required for oncogenic progression in pancreatic cancer.

Furthermore, deletion of MBL or C3 in the extra-tumoral compartment or knockdown of C3ar in tumor cells were both protective against tumor growth in mouse models. There was thus a suggestion that using anti-fungal agents at some point of the treatment of pancreatic cancer may result in a shrinking or slowing down the growth of the tumor. In a paper reviewing expression of complement in various cancers most cancers over-expressed C3 but neutrally-expressed C5, and the most over expressed gene was CD59, suggesting efficient protection of malignant cells from complement-mediated killing (9).

To document the expression of C3 and *Malassezia* and associated genes in our patient population, we decided to perform studies on paraffin embedded pancreatic cancer biopsies using pancreatic cancer and their normal tissue counterpart samples from the University of New Mexico Tissue Repository.

## Results

### Microbiome results

Using optimized methods (10), we used RNA-seq analysis on paired tumor-normal PDAC tissue samples derived from formalin-fixed paraffin embedded (FFPE) slices for 15 patients obtained from University of New Mexico Tissue Repository, generating high quality data for 14 patients, with an average of 32 × 10^-6 reads per sample (Table 1, Table S1). After sequencing, reads were taxonomically classified using kraken2 (11, 12, 13) and braken (14) against a library containing human, viral, bacterial, and fungal genomes, including the genomes of several *Malassezia* species. The fungus *Malassezia* was present in all tumor samples at varying concentrations (Table1, TableS1).

**Table 1:** RNA-sequening statistics

| Table 1: RNA-sequencing statistics |  |  |  |  |  |
| --- | --- | --- | --- | --- | --- |
| | No. Samples | Average Total Reads, $\times 10^6$ (range) | Average Human Reads, $\times 10^6$ (range) | Mean Number of Genera (range) | Mean Percent Reads *Malassezia* (range) |
| Normal Samples | 14 | 27.0 (14.4 - 40.0) | 13.0 (6.9 - 19.4) | 220 | 0.04 |
| Tumor Samples | 14 | 36.6 (12.2 - 96.4) | 17.7 (5.9 - 46.6) | 224 | 0.02 |

We also detected *Malassezia* in all of the normal tissue samples, which is in contrast to a previous study (Table1, TableS1) (8). Despite reports that PDAC tumors have a lower microbial diversity than normal pancreatic tissues (6), we found no significant difference in the number of microbial genera present in normal samples versus tumor samples (Table 1)

### Transcriptome Analysis

Unsupervised multidimensional scaling (MDS) was used to compare the normal and tumor samples. The MDS plot (Figure 1A) revealed two tumor sample outliers, which were more similar to the normal samples than the other tumor samples. In addition, most of the normals clustered tightly together, while the tumor samples were more dispersed, indicating that the tumor gene expression pattern were more heterogenous while the normals were more homogenous. Differential gene expression (DGE) analysis of the 14 paired tumor-normal patient samples was used to characterize transcriptome differences between tumor and normal samples. As shown in Figure 1B, we found 3,177 genes that were at least 1.5-fold up or down regulated (up = 1593, down = 1584) and had an adjusted p-value less than or equal to 0.05. The heatmap in Figure 1C summarizes the dramatic differences in gene expression in normal and tumor samples.

**Figure 1:**
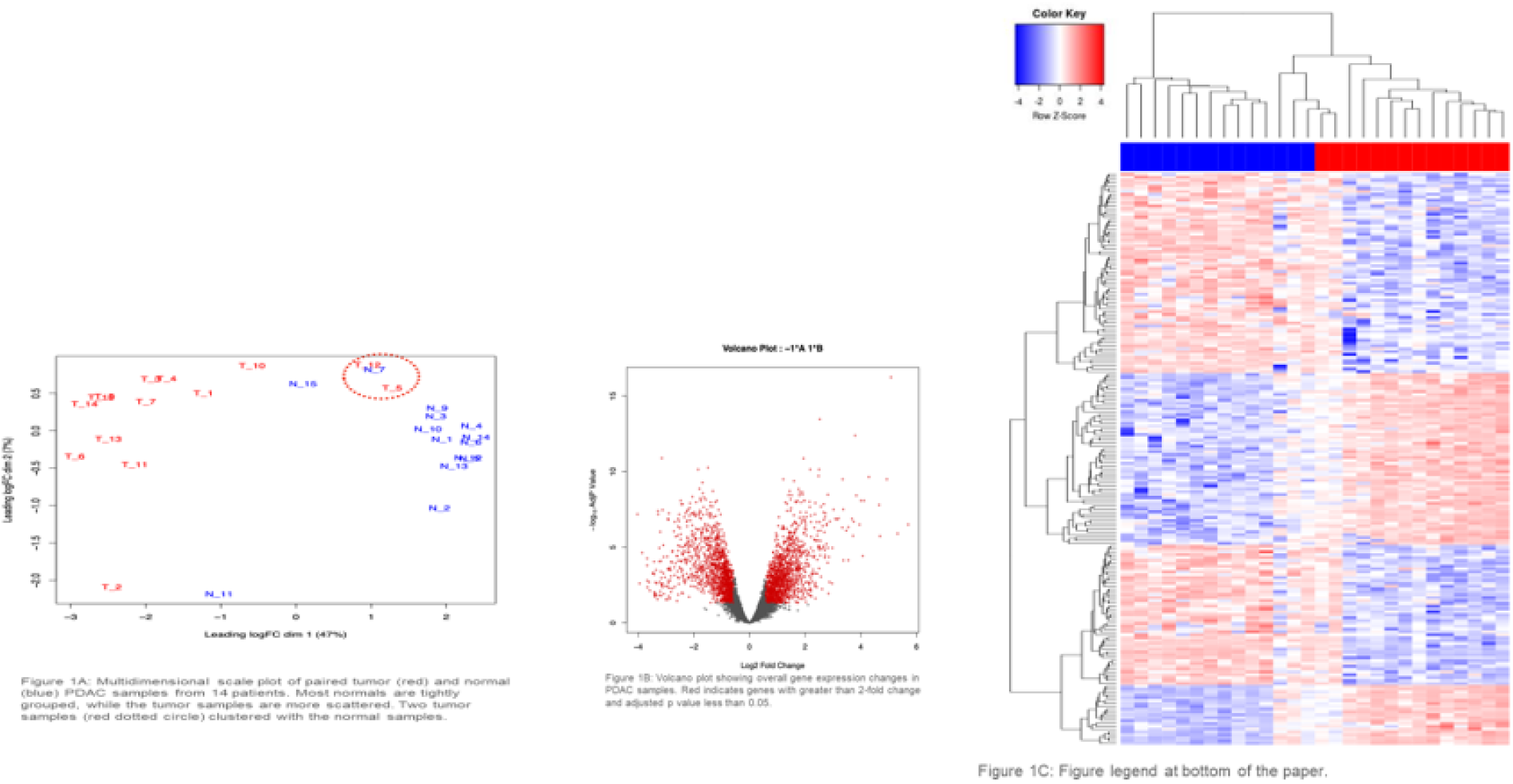
(**A**) Multidimensional scale plot of PDAC tumor (red bar) and normal (blue bar) RNA-seq data. (B) Volcano plot summarizing the differential gene expression analysis, showing log2 fold change vs. log10 of the *p*-value (BH adjusted). Red indicates genes that were >= 1.5 log2 fold change and had an adjusted p-value <= 0.05. (**C**) The heatmap summarizes the supervised clustering and differential gene expression analysis comparing the normal samples to the tumor samples. Dendrograms show the hierarchical clustering of samples (top) and genes (left side). A larger version of this heatmap with samples and genes labeled is provided in the supplementary results (Supplementary Figure 1).

*Malassezia* is proposed to promote tumor growth via the complement pathway, disruption of which activates inflammation (15, 8). Gene set enrichment analysis (GSEA) can reveal pathway and disease phenotypes within sets of differentially expressed genes. GSEA of the set of genes differentially expressed between tumor and normal samples against the curated (C2), ontology (C5), and hallmark (H) gene sets at the molecular signatures database (16,17), revealed enrichment for genes involved in the complement cascade, complement activation, and inflammatory response. Additionally, our differentially expressed genes are enriched for expected gene sets such as pancreatic cancer, epithelial mesenchymal transition, and KRAS signaling (Table 2).

**Table 2:**
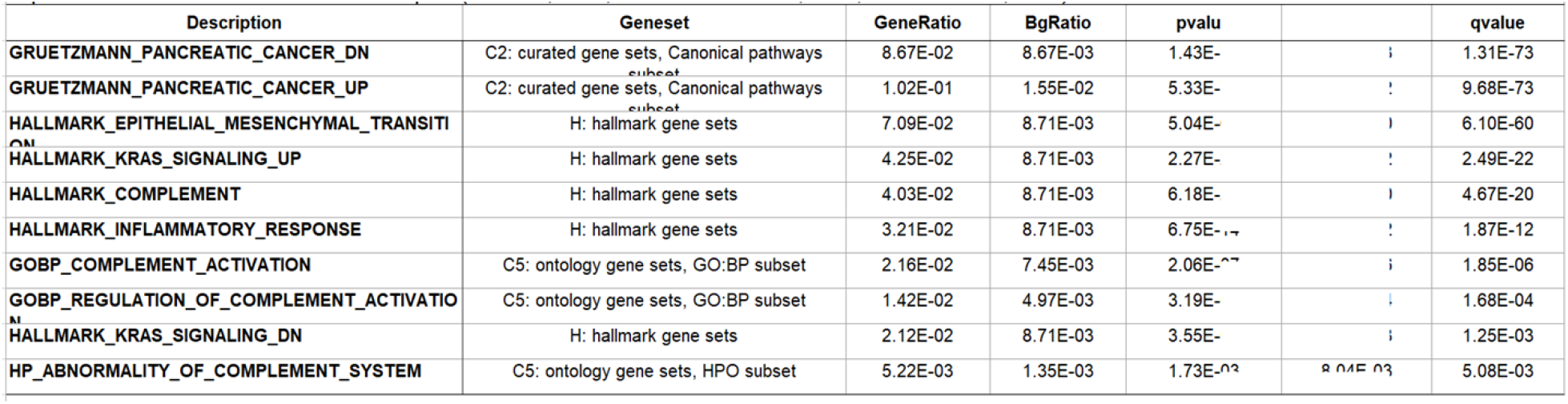
Select gene set enrichment analysis results. Clusterprofiler was used to probe genesets at the Molecular Signatures Database using genes differentially expressed between tumor and normal samples (Yu et al., 2012, Subramanina et al., 2005, Liberzon et al., 2015).

There is an extensive literature of the linkage between the complement/inflammation genes and cancer, including PDAC, focusing primarily on the tumor microenvironment and how immune cells can affect tumor growth through the inflammatory process (18, 19). Since each of these pathways were significantly enriched in our dataset (Table 2), we specifically examined which genes were differentially expressed, and their directionality. As expected, only a subset of the genes in each gene set were significantly differentially regulated: 64 of 200 for complement (Figure 2A), 52 of 200 for inflammatory (Figure 2B), and 104 of 200 for EMT (Figure 2C).

**Figure 2:**
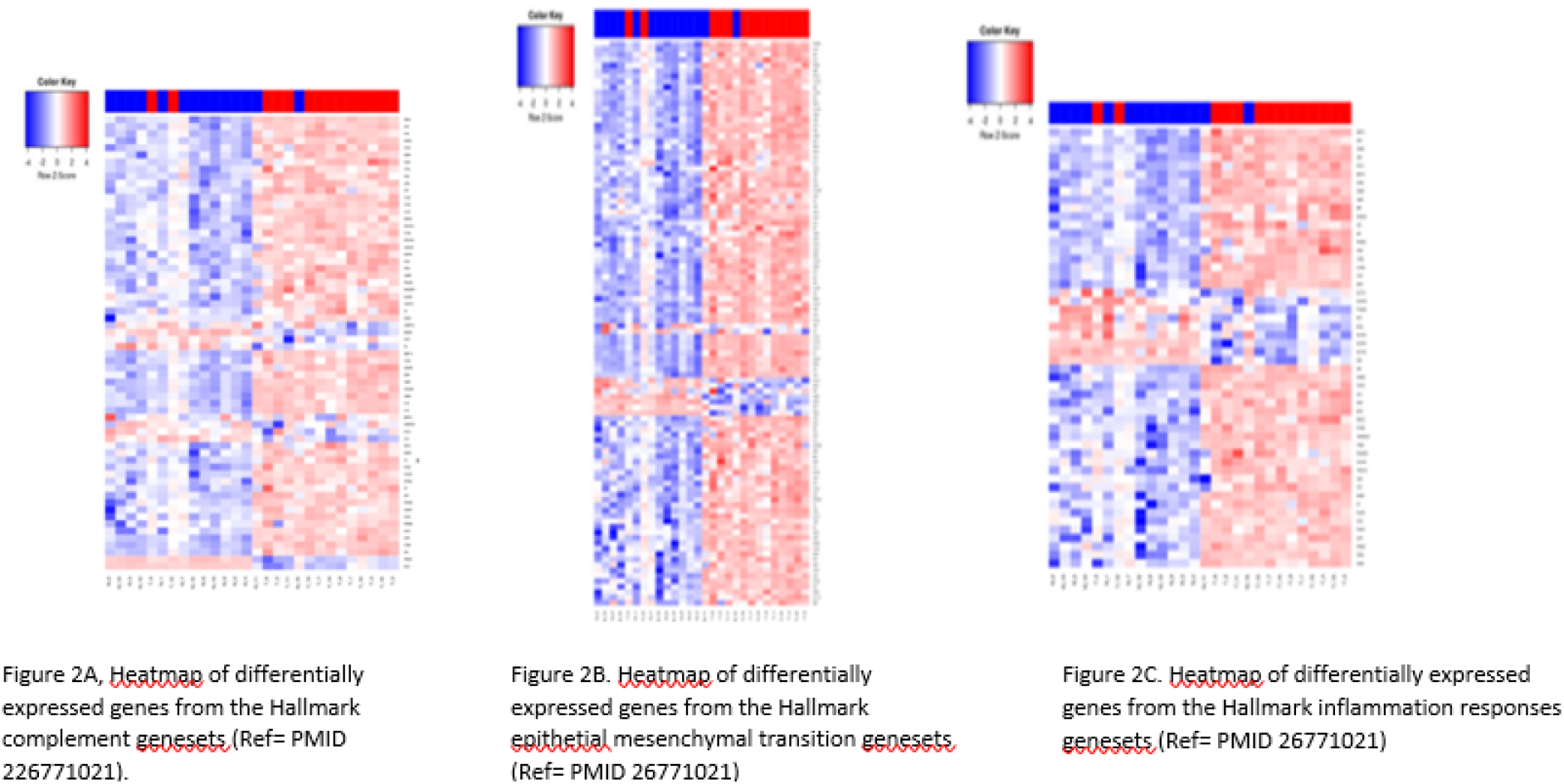
Heatmaps of geneset genes that are differentially expressed between tumor (red bar) and normal (blue bar) PDAC samples. (A) Hallmark Complement Geneset, the C2 gene is indicated by (*). (B) Hallmark Epithelial Mesenchymal Transition Geneset. (C) Hallmark Inflammation Response Geneset. (Ref = PMID 26771021).

Interestingly, overall expression of C3 was higher than C2 expression on both tumors and normals, but only C2 was significantly up-regulated in tumors (Figure 3). C2 is a part of the lectin and classical complement pathways which result in the activation of C5 and the membrane attack complex (MAC). C3dg is a cleavage product of the C3 component of complement that can facilitate the co-ligation of the complement receptor 2 (CR2/CD21) with the BCR via C3dg/Ag complexes which can amplify BCR-mediated signaling events and lower the threshold for B cell activation. They also investigated CR2-mediated stimulation of peripheral B cell subpopulations and demonstrated that CR2 is capable of amplifying BCR signal transduction in each subpopulation of B-2 cells, but not B-1 cells. (20).

**Figure 3:**
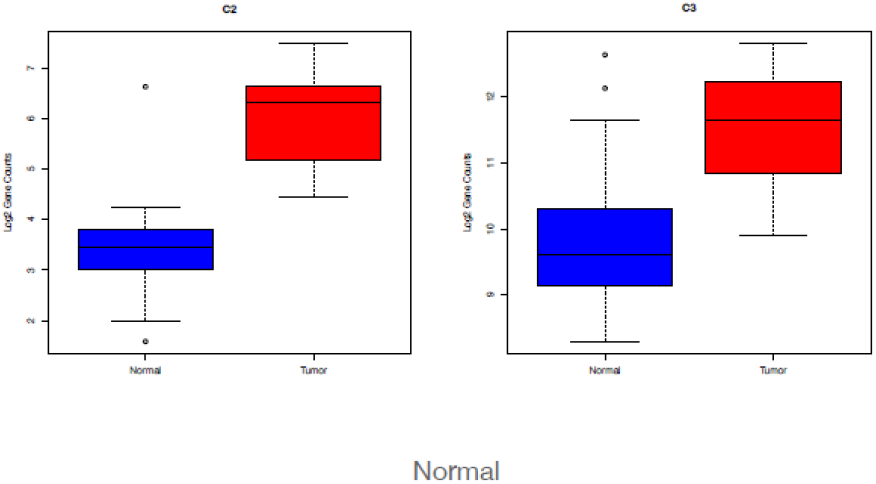
Boxplots of log2 gene counts for C2 (left) and C3 (right) in normal samples (blue) and tumor samples (red).

We were particularly interested in the expression pattern of genes known to play a role in pancreatic cancer, complement, inflammation and EMT. Using the genesets described above, as well as a pancreatic cancer genesets (Gruetzmann pancreatic cancer up and Gruetzmann pancreatic cancer down, (21) we looked for genes involve in multiple processes that indicate networks of influential genes using a CNET plot (Figure 4). This method showed that there were very few genes that overlapped the down regulated pancreatic cancer genes list and the complement pathway (1 gene), inflammatory response (1 gene), and EMT (3 genes) lists (Figure 4). However, we observed a number of genes that were up-regulated in pancreatic cancer and involved in both EMT (229 genes) and the complement pathways (14 genes), but a relative small number of inflammatory response genes (6 genes), all of which were up-regulated in our tumor samples. Interesting, PLAUR (urokinase plasminogen activator receptor) overlapped with all of these pathways, while FN1 (fibronectin 1) overlapped with both complement and EMT processes.

**Figure 4:**
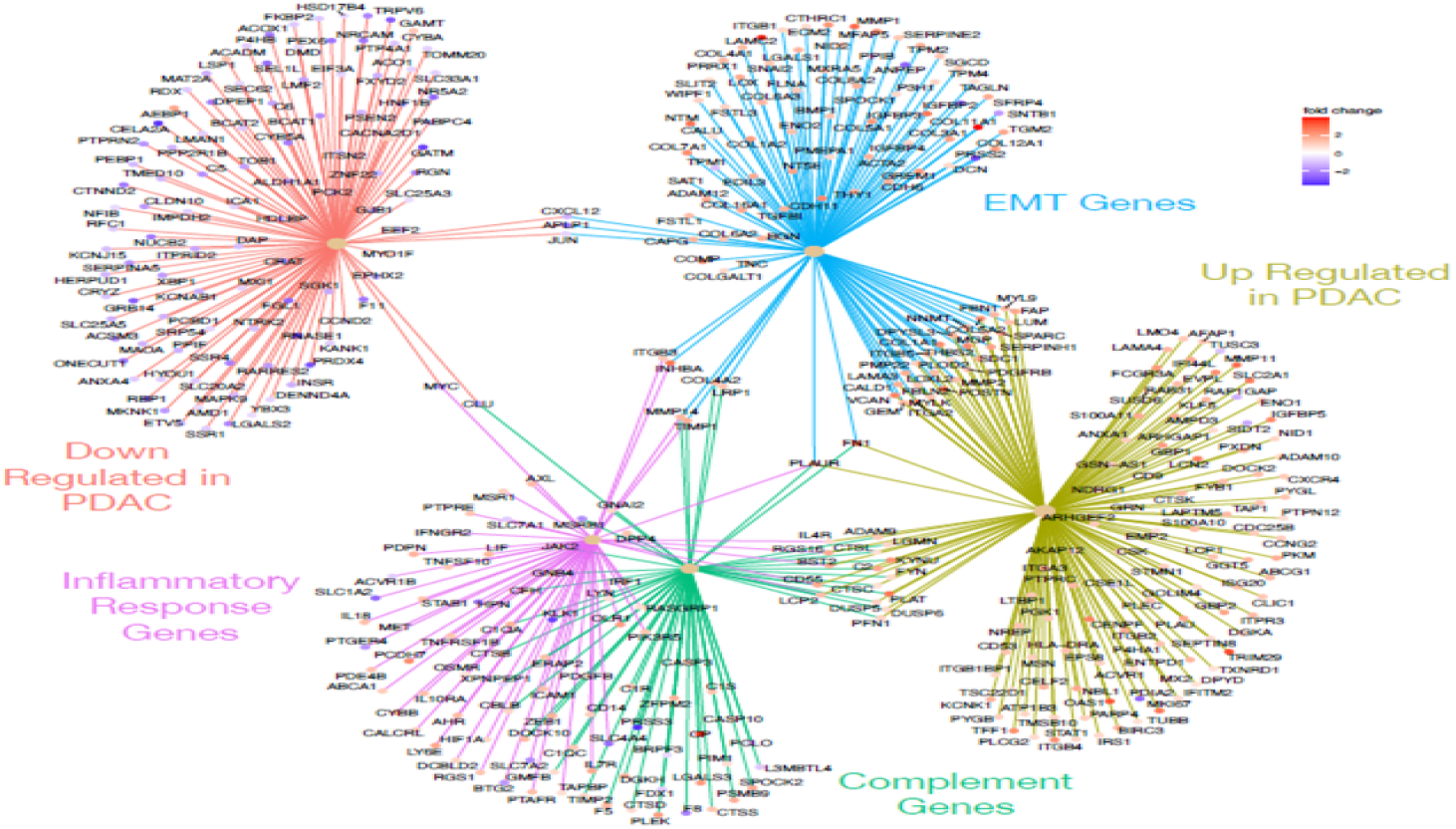
CNET plot showing gene overlaps in enriched genesets. Geneset enrichment analysis of genes differentially expressed between tumor and normal samples indicated enrichment for sets of genes up (olive) or down (orange) regulated in pancreatic cancer, the complement cascade (green), epithelial mesenchymal transition (blue), and inflammatory response (purple). Genes at the end spokes indicate differential expressed in our dataset that were enriched in each geneset, color of dots indicated whether the gene was up (red tones) or down (blue tones) regulated in tumors relative to normal samples. (17, 21, 34)

In a recent paper another group established that Kras G12D regulates the expression of a chemo attracting cytokine, IL-33, that recruitsTH2 and innate lymphoid cells (ILC2 cells). The TH2 and ILC2 cells via their pro-tumorigenic cytokine production accelerate PDAC tumor progression. They established IL-33 as a downstream target of Kras G12D and that IL-33 expression is significantly upregulated in PDAC patients. They also identified the role for the intratumoral mycobiome (*Malassezia* and *Altermata*) in facilitating the type 2 immune response in the PDAC TME by stimulating the extra cellular secretion of IL-33. The role of intra-tumor fungus is distinct from prior observation whereby fungi were shown to activate a complement system (8) and might act in tandem to promote PDAC progression. These mechanistic insights governing the extracellular secretion of IL-33 suggested that the IL-33-ILC2/TH2 axis may be a potential target in PDAC. They found that fungal-derived components can activate a dectin-1 mediated Src-Syk-CARD9 pathway which, via downstream mechanisms, secretes IL-33. Genetic deletion of IL-33 or anti-fungal treatment caused PDAC tumor regression. Based on these findings, they concluded that intra-tumoral fungi or fungal products provoke IL-33 secretion by PDAC cells, which promotes type 2 immune responses and tumor progression in this PDAC subset (22).

Since our differentially expressed genes were also enriched for the Hallmark KRAS signaling pathway (Table 2), we also looked for overlaps between the KRAS signaling pathway geneset, and the other three sets used above. Figure 5 shows the CNET plot for the genesets alone, but the intersection of the geneset plus the PDAC genesets can be seen in Figure S2. Our differentially expressed genes contain 5 genes which overlap both KRAS and complement pathways (PLAT, CFH, CTSS, DUSP6, and PLAUR, Figure 5), all of which are up-regulated in tumors. Interestingly, PLAUR (urokinase plasminogen activator (uPA) and/or its receptor (uPAR)) overlaps with all of the tested geneset (PDAC genes, complement and KRAS pathways, inflammatory and EMT genes, Figures 4-5, Figure S2), suggesting this is a good drug target for the treatment of PDAC. Urokinase plasminogen activator (uPA) and/or its receptor (uPAR) are essential for metastasis, and overexpression of these molecules is strongly correlated with poor prognosis in a variety of malignant tumors. Notably impairment of uPA and/or uPAR function impedes the metastatic potential of many tumors (23).

**Figure 5:**
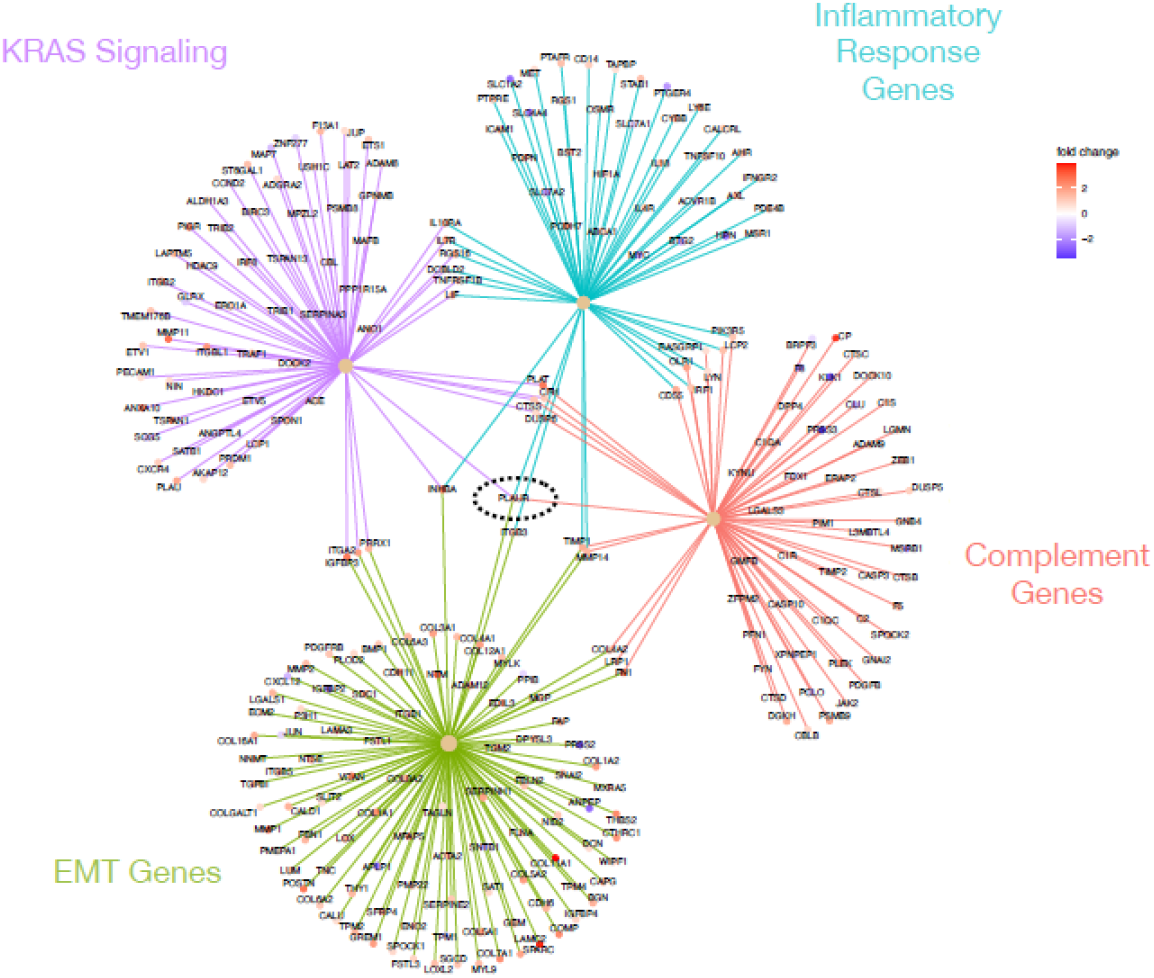
CNET plot showing gene overlaps in enriched genesets. Geneset enrichment analysis of genes differentially expressed between tumor and normal samples indicated enrichment for sets of genes involved in KRAS signaling (purple), inflammatory response (turquoise), complement cascade (red), and epithelial mesenchymal transition (green). Black dashed circle indicates PLAUR, which connects to all four genesets. Genes at the end spokes indicate differential expressed in our dataset that were enriched in each geneset, color of dots indicated whether the gene was up (red tones) or down (blue tones) regulated in tumors relative to normal samples. (17, 21 34).

We also found over-expression of Gal-3 and Dectin-1 in the tumor samples compared to the normal samples (figure 6). Galectins are a group of evolutionarily conserved proteins with the ability to bind β-galactosides through characteristic carbohydrate-recognition domains (CRD). Galectin-3 is structurally unique as it contains a C-terminal CRD linked to an N-terminal protein-binding domain, being the only chimeric galectin (24). We found over-expression of Gal-3 in the tumor samples compared to the normal samples. This has been replicated in other studies. Two studies demonstrated that Gal-3 is highly up-regulated in pancreatic tumor tissues and cells in both human pancreas and in a K-ras mutant mouse model of pancreatic cancer. The data suggest that Gal-3 in tumor tissues further enhances Ras activity by binding and retaining Ras at the plasma membrane, thereby activating down-stream Ras signaling. Gal-3 may thus play a role in the pathogenesis of PDAC, in which Ras mutations frequently occur (25, 26).

**Figure 6:**
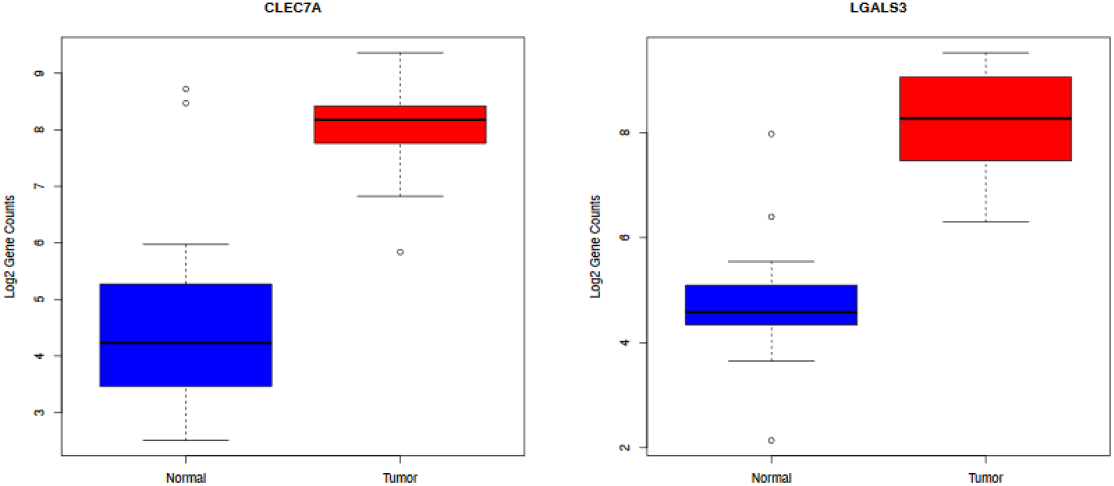
Boxplots of log2 gene counts for CLEC7A (DECTIN-1 (left)) and LGALS3 (Galectin-3 (right)) in normal samples (blue) and tumor samples (red).

Dectin-1 recognizes β-glucans with its carbohydrate recognition domains (CRD) and transduces signals through its immune-receptor tyrosine-based activation motif (ITAM)-like motif in the cytoplasmic domain. β-glucans is a major cell wall component of fungi (27).

A recent study found that Dectin-1 is highly expressed on macrophages in pancreatic ductal adenocarcinoma (PDAC). Dectin-1 signaling resulted in PDAC progression, while Dectin-1 deletion or blockade of its downstream signaling decreased tumor growth. They showed that Syk phosphorylation was a downstream signal for Dectin-1 (28). Furthermore the recent paper describing the effect of the fungus on PDAC subset (Kras G12D) via an alternative pathway (the IL-33-ILC2/TH2 axis) is activated by a dectin-1 mediated Src-Syk-CARD9 pathway (22).

## Discussion

We explored the TME in the samples and found overexpression of C2, Dectin-1, and Galectin-3 represented in box plot analysis. There was no literature about the role of C2 in pancreatic cancer, but there was data showing C2 has a role in amplifying BCR signal transduction in each subpopulation of B-2 cells, but not B-1 cells. Dectin-1 and Galectin-3 can augment PDAC progression as noted above. These two molecules can propagate the growth of the PDAC in vivo and mice models, but this must be looked at in the context of a multi molecular array, some inhibiting and some promoting PDAC growth.

We also compared the gene expression profiles of PDAC patient tumor samples to their paired normal sample and found several differential expressed genes. These genes were enriched for genes that are known to be involved in the complement cascade and inflammation, as well as genes involved in epithelial-mesenchyme transition, KRAS signaling, and known PDAC genes. From this we were able to identify one gene (PLAUR) that sits at the intersection of these pathways and may be a potential target for PDAC treatment. In addition, we found a handful of genes that represent additional attractive drug targets because they overlap subsets of complement, inflammation, KRAS and EMT pathways. Additionally, these results suggest that the TME contributes to PDAC progression by activation of these critical pathways.

We did not have the data on the clinical outcome of the patients. Thus no biological correlative analysis was performed. Unlike the Aykut B et.al (8) and Alam A et al (22) groups we noted the fungus *Malassezia* in the pancreatic cancer and normal control in relatively the same proportions. However, despite the fact that there was no difference in *Malassezia* in tumors versus normal tissue, there was a significant difference in expression of a number of genes related to inflammation, infection and EMT as noted in Figure 6. The role of the microbiome, including fungi is being recognized as having a significant role in the pathophysiology and progression of PDAC, and we believe solving how to alter the flora around and in the tumor can add another dimension to treating this disease. In summary, we have demonstrated the interaction between proteins involved in inflammation can also have a role in propagation of growth in pancreatic cancer growth.

## Methods

### RNA Isolation and Sequencing

Total RNA was extracted from FFPE slices using the RNeasy FFPE kit (Qiagen) and the manufacture’s protocol. Synthesis of cDNA and library preparation were performed using the SMARTer Universal Low Input RNA Kit for Sequencing (Clontech) and the Ion Plus Fragment Library Kit (ThermoFisher) as previously described (29, 10, 30). Sequencing was performed using the Ion Proton S5/XL systems (ThermoFisher) in the Analytical and Translational Genomics Shared Resource at the University of New Mexico Comprehensive Cancer Center.

RNA sequencing data is available for download from the NCBI BioProject database using study accession number PRJNA940178.

### Data Analysis

Prior to alignment, non-human RNA-seq reads were identified and removed from analysis using the kraken2 taxonomic sequence classification system against a library containing human, fungal, bacterial and viral genomes (11,12,13) and final genera level abundances were calculated using Bracken (v2.5.0, (14). Following kraken2 filtering, all sequences classified as human were aligned using tmap (v5.10.11) to a BED file containing non-overlapping exons from UCSC genome hg38. Exon counts were calculated using HTseq (v0.11.1, (31), and gene counts were generated by summing counts across exons. Samples were normalized for library size using edgeR (32) and low expressing genes were excluded from the final analysis using a filtering threshold of 20 reads in 9 samples. EdgeR and DESeq (33) were used for principal component analysis. EdgeR was also used for the differential expression (crosswise comparison of the groups) using the glm method with an adjusted p-value of cutoff of 0.05 and requiring a minimum fold change of 1.5. Differentially expressed genes were further analyzed using various R packages including clusterProfiler (34). For additional details see (29) and (10).

## Supplemental Figures

**Figure S1:**
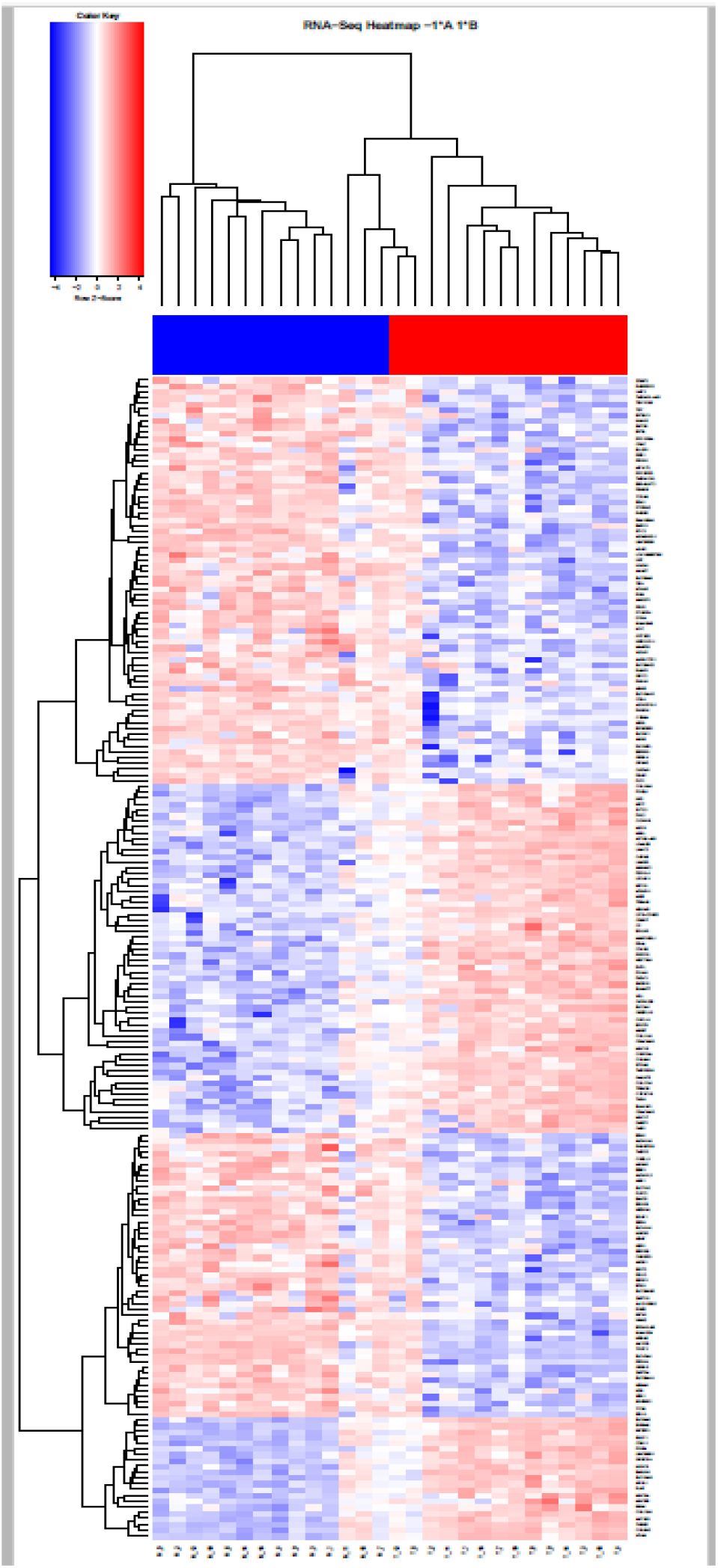
Heatmap summarizing the supervised clustering and differential gene expression analysis comparing the normal samples to the tumor samples. Sample names are along the bottom, gene names are along the right side of the figure.

**Figure S2:**
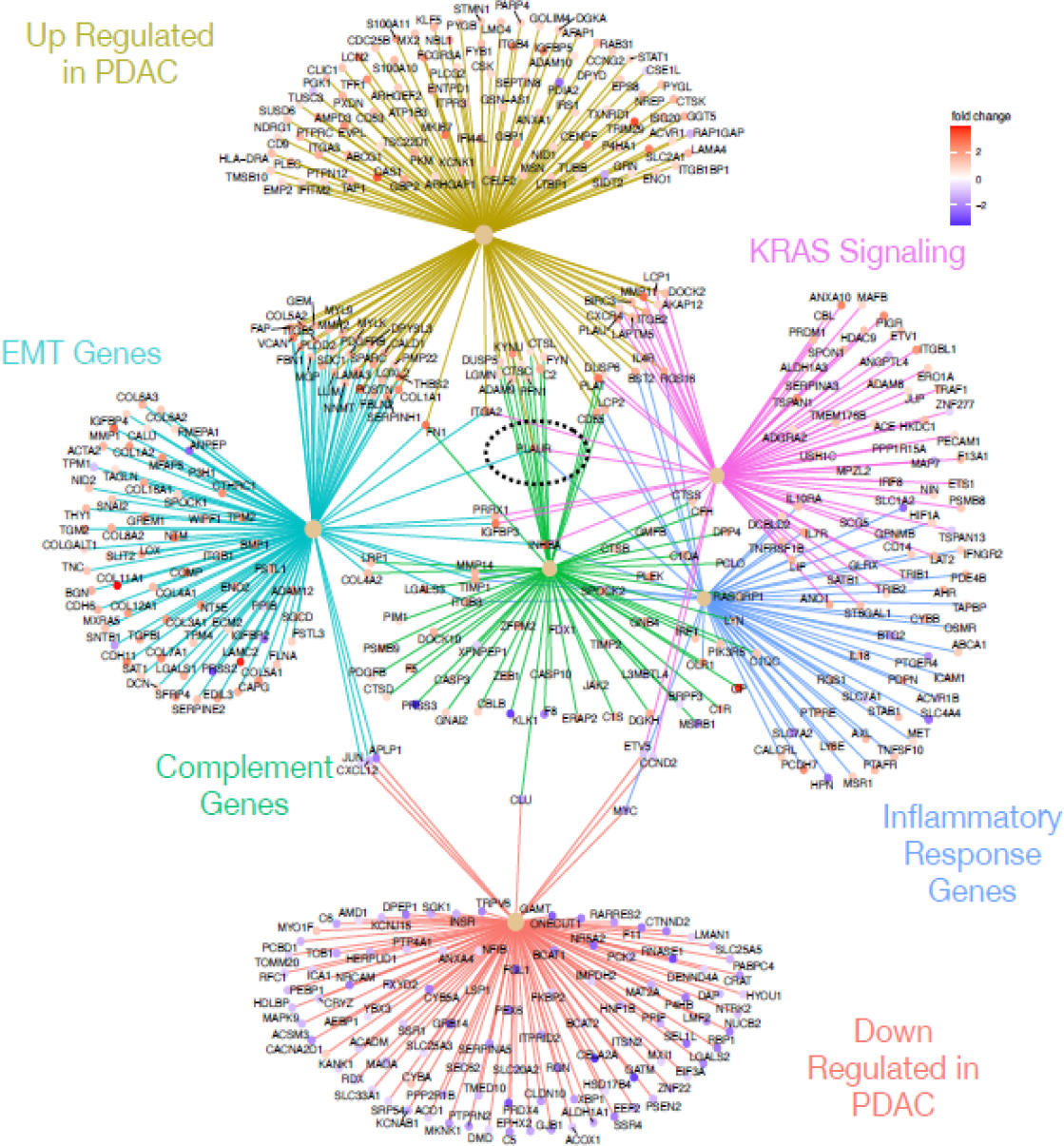
CNET plot showing gene overlaps in enriched genesets. Geneset enrichment analysis of genes differentially expressed between tumor and normal samples indicated enrichment for sets of genes involved in KRAS signaling (purple), inflammatory response (blue), complement cascade (green), and epithelial mesenchymal transition (turquoise). Figure also includes genes known to up (orange) or down (olive) regulated in PDAC. Black dashed circle indicates PLAUR, which connects to all 5 of the 6 genesets (not connected to down in PDAC). Genes at the end spokes indicate differential expressed in our dataset that were enriched in each geneset, color of dots indicated whether the gene was up (red tones) or down (blue tones) regulated in tumors relative to normal samples. (17, 21, 34).

**Table S1.**
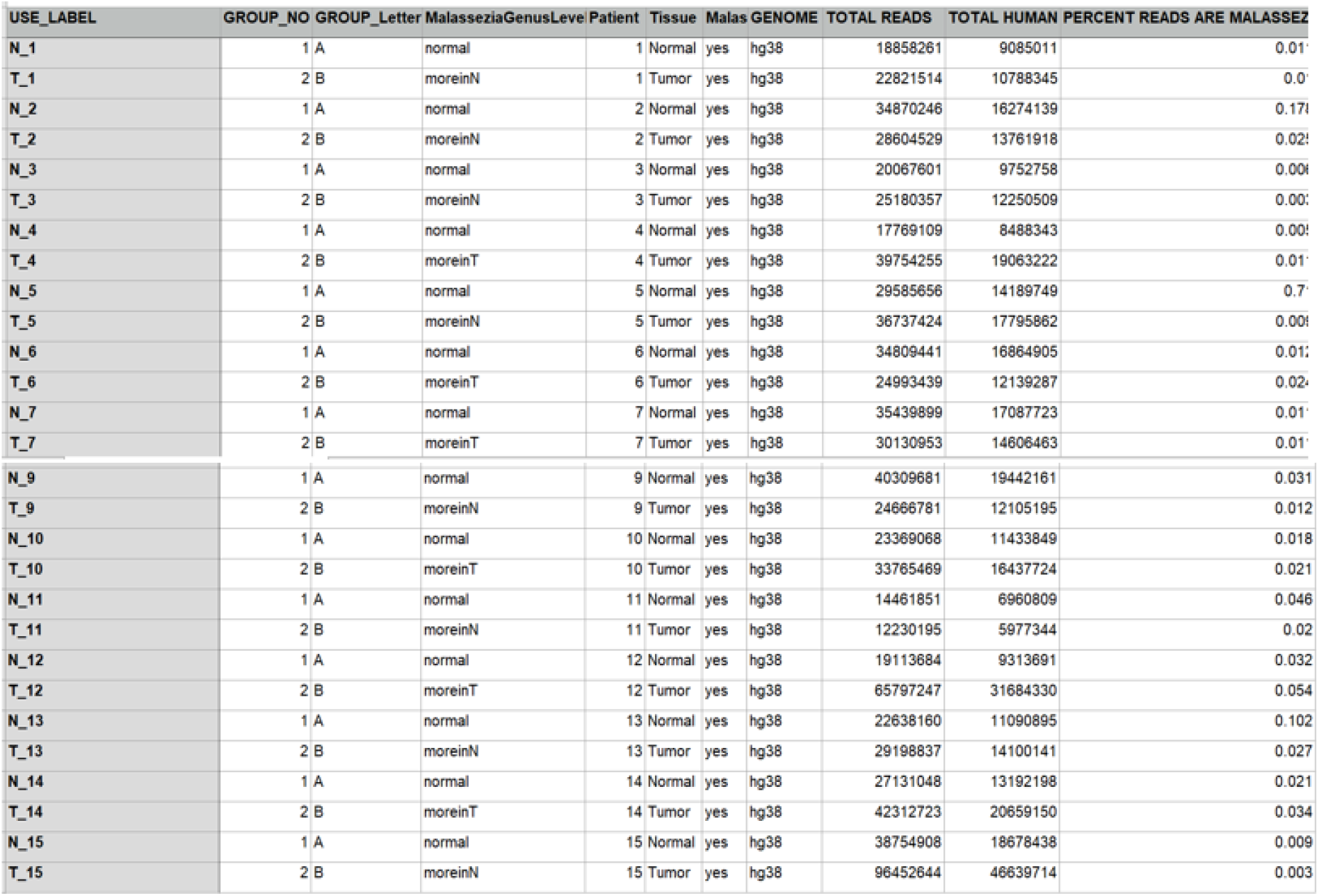
RNA sequencing statistics by sample.

